# Perceptual learning evidence for an interval- and modality-invariant representation of subsecond time

**DOI:** 10.1101/794032

**Authors:** Ying-Zi Xiong, Shu-Chen Guan, Cong Yu

**Affiliations:** Psychology, McGovern Brain Institute, and Center for Life Sciences, Peking University

## Abstract

A central theme in time perception research is whether subsecond timing relies on a dedicated centralized clock, or on distributed neural temporal dynamics. A fundamental constraint is the interval- and modality-specificity in perceptual learning of temporal interval discrimination (TID), which argues against a dedicated centralized clock, but is more consistent with multiple distributed mechanisms. Here we demonstrated an abstract, interval- and modality-invariant, representation of subsecond time in the brain. Participants practiced TID at a specific interval (100 ms), and received exposure to a transfer interval (200 ms), or to a different auditory/visual modality, through training of an orthogonal task. This double training enabled complete transfer of TID learning to the untrained interval, and mutual complete transfer between visual and auditory modalities. These results demonstrate an interval- and modality-invariant representation of subsecond time, which resembles a centralized clock, on top of the known distributed timing mechanisms and their readout and integration.

## Introduction

Accurate timing is critical in sensory and cognitive tasks, especially those requiring subsecond precision, such as understanding conversations and listening to music. Subsecond timing is first thought to be based on a dedicated centralized clock, which acts like a pacemaker and an accumulator to keep track the time (Creelman, 1962; Treisman, 1963). Later studies found that timing is intrinsic properties of neural dynamics that distribute over many sensory modalities and brain areas (Ivry & Schlerf, 2008; Paton & Buonomano, 2018). Temporal ramps in neuronal firing (Durstewitz, 2003), peaks in sequential activations of neuron pools (Hass et al., 2008), temporal evolution of the collective state of an entire neural network (Buonomano, 2000), and cortical oscillations at various frequency bands (Matell & Meck, 2004; Allman et al., 2014), all carry information sufficient for subsecond timing (Hass & Durstewitz, 2016). However, centralized and distributed timing models do not necessarily completely contradict to each other, as distributed neural temporal information needs to be readout and integrated by more centralized decision mechanisms for timing perception (Shi et al., 2013; Paton & Buonomano, 2018).

Regardless, almost all models of subsecond timing are constrained by the frequent observations that TID perceptual learning is always interval specific. Wright et al. (1997) first reported this specificity in that auditory TID learning at a 100-ms interval cannot transfer to a 50-ms or 200-ms interval at the same tone frequency. The interval specificity has been replicated by various sensory and motor TID learning studies (Nagarajan et al., 1998; Meegan et al., 2000; Karmarkar & Buonomano, 2003; Wright et al., 2010). These observations appear to be inconsistent with models of a centralized clock (Creelman, 1962; Treisman, 1963) that would not predict interval specificity. Rather they are more consistent with distributed mechanisms of intrinsic neural timing, especially with evidence that cortical slices *in vitro* can learn a specific interval after repeated presentations (Goel & Buonomano, 2016). They may also indicate separate readout units, each assigning more weights to inputs representing a specific interval from distributed mechanisms (Bueti & Buonomano, 2014; Paton & Buonomano, 2018). Alternatively, the interval specificity can be interpreted as indications of multiple centralized clocks, with each responsible for a specific time interval (Ivry & Richardson, 2002).

In addition, timing information from various distributed mechanisms have to be integrated, probably on the basis of Bayesian priors for optimal decision making (Cicchini et al., 2012; Shi et al., 2013). For example, more precise auditory timing information is given more weight than less precise visual timing information. Consistent with this Bayesian process is the observations that auditory TID learning can transfer to visual TID, but visual TID learning cannot transfer to auditory TID learning, suggesting the dominance of auditory timing in decision making (Bratzke et al., 2012; McGovern et al., 2016).

However, when perceptual learning does not show transfer, it does not necessarily mean that learning is specific. That is, the transfer of learning could be hindered by some other factors associated with conventional training (i.e., training one condition and pre- and post-testing an untrained condition). Inspired by our previous perceptual learning studies, we suspected that TID learning is actually completely transferrable across intervals, as well as across visual and auditory modalities. To test these possibilities, we applied a double-training procedure that we originally developed for visual perceptual learning research (Xiao et al., 2008; Zhang et al., 2010; Xiong et al., 2016). Double training consists of two training tasks. The primary training task here was temporal interval discrimination. The secondary training task was an orthogonal one at the untrained interval or modality (see Results for details). The secondary training task was “orthogonal” because it did not affect the performance of the primary TID task by itself. Rather its purpose was to activate sensory neurons responding to the untrained interval or modality, so that the potentially interval- and modality-invariant TID learning can now functionally connect to the new inputs to enable complete learning.

We have successfully made perceptual learning of various visual tasks transfer significantly, and often completely, to new retinal locations or orientations with double training (Xiao et al., 2008; Zhang et al., 2010; Wang et al., 2013; Wang et al., 2014; Xiong et al., 2016), as well as visuomotor learning to new directions (Yin et al., 2016) and auditory tone frequency learning to new frequencies (Xiong et al., 2019). If double training could also make TID learning transfer to new intervals and across visual/auditory modalities, we would have evidence for an abstract interval- and modality-invariant representation of subsecond time, which resembles the idea of a centralized clock in terms of centralized and interval-invariant processing of time. This would imply that the centralized and distributed mechanism models of subsecond timing are actually complementary, each addressing different aspects of the complex subsecond timing processes.

## Results

### Baselines: Interval and auditory/visual modality specificity in TID learning

We first replicated the interval specificity of TID learning (Wright et al., 1997; Nagarajan et al., 1998; Meegan et al., 2000; Karmarkar & Buonomano, 2003; Wright et al., 2010) and the asymmetric learning transfer from auditory to visual TID (Bratzke et al., 2012; McGovern et al., 2016) to establish the baselines for later experiments. One group (N = 8) practiced auditory TID at a 100-ms standard interval (auditory TID training baseline group), and a second group (N = 9) practiced visual TID at the same 100-ms standard interval (visual TID training baseline group).

For the auditory TID training group, training reduced auditory TID thresholds at a 100-ms interval by 0.29 ± 0.07 log units (t = 5.41, p < 0.001, Cohen’s d = 1.91, 95% CI [0.18, 0.39]; see Methods for details of data analysis). But it had no significant impact on auditory TID at a 200-ms interval, reducing thresholds by merely 0.08 ± 0.04 log units (t = 1.47, p = 0.15, Cohen’s d = 0.52, 95% CI [-0.03, 0.18]), replicating the known interval specificity (Fig. 1a, b).

**Figure 1.**
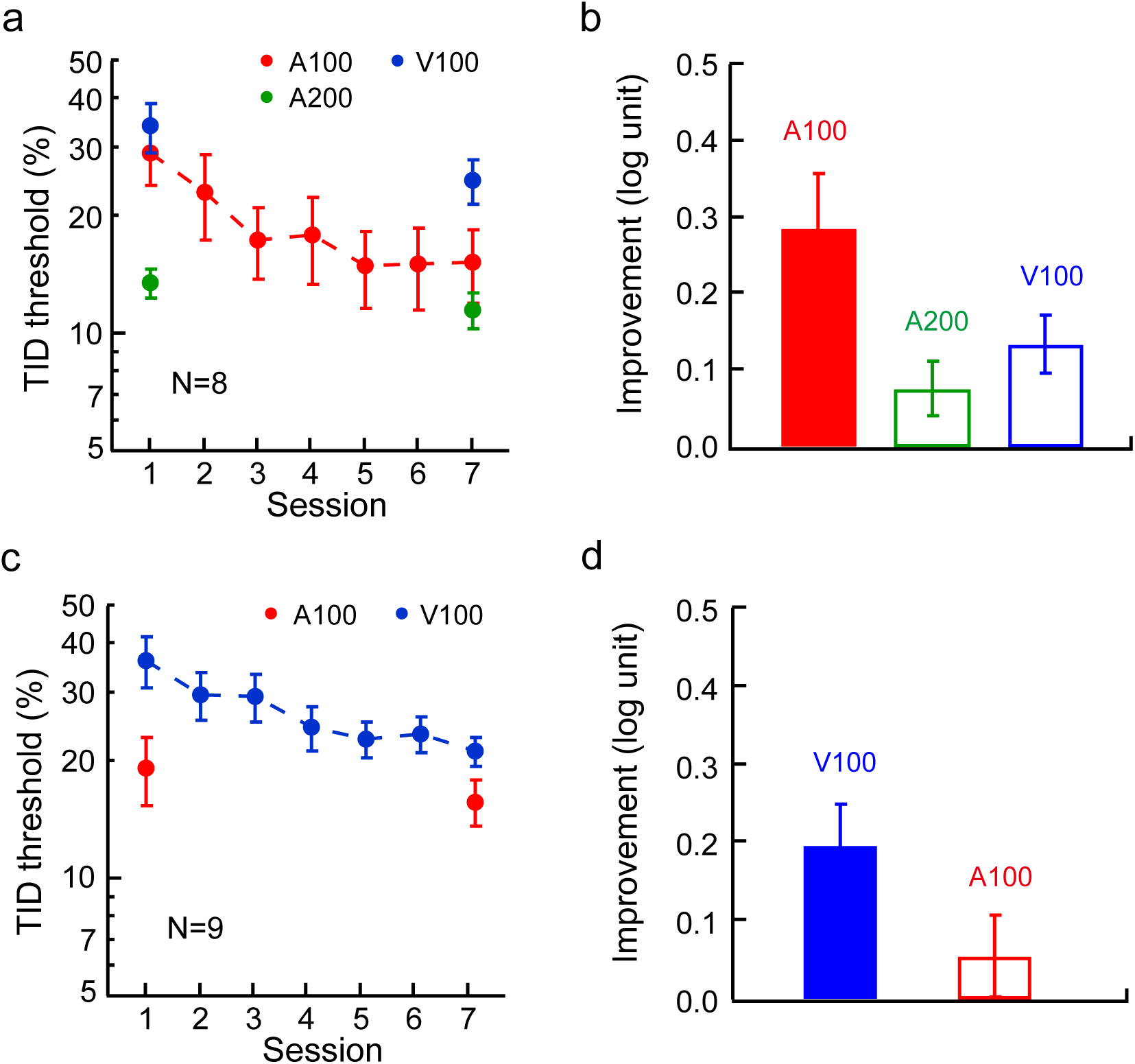
Baselines: Interval specificity and asymmetric auditory-to-visual transfer in temporal interval discrimination (TID) learning. **a**. The mean auditory TID learning curve at a 100-ms interval (A100), and pre- and post-training auditory TID thresholds at a 200-ms interval (A200) and visual TID thresholds at a 100-ms interval (V100). **b**. The mean improvements of TID with trained A100, untrained A200, and untrained V100 conditions. **c**. The mean visual TID learning curve at a 100-ms interval (V100), and the pre- and post-training auditory TID thresholds the same interval (A100). **d**. The mean improvements of TID with trained V100 and untrained A100. Here and in later figures, error bars indicate ±1 standard error of the mean, and solid and empty histograms indicate threshold improvements from training and transfer, respectively.

The auditory TID training also improved visual TID at the same 100-ms interval, reducing visual TID thresholds by 0.14 ± 0.04 log units (t = 2.56, p = 0.012, Cohen’s d = 0.90, 95% CI [0.03, 0.24]) (Fig. 1a, b). However, for the visual TID training baseline group, although training improved visual TID at a 100-ms interval (by 0.20 ± 0.05 log units; t = 3.96, p < 0.001, Cohen’s d = 1.32, 95% CI [0.10, 0.30]), the learning did not transfer to auditory TID at the same interval (by 0.05 ± 0.05 log units; t = 1.08, p = 0.28, Cohen’s d = 0.36, 95% CI [-0.05, 0.15]) (Fig. 1c, d). These data thus replicated the asymmetric transfer of TID learning from auditory to visual, but not the other way around. Here the visual TID improvement through auditory TID training (V100 in Fig. 1b) was about 70% of that through direct visual TID training (V100 in Fig. 1d), suggesting that auditory TID training might have not maximized the visual TID performance in these observers.

### Complete transfer of TID learning across intervals with double training

We then investigated whether double training could abolish interval specificity, a major constraint on modeling the mechanisms of subsecond time perception. Eight participants practiced auditory TID at a 100-ms interval, as well as an orthogonal tone frequency discrimination task at a 200-ms interval, in alternating blocks of trials for five sessions. Our hypothesis was that this secondary frequency discrimination training would allow the participants to receive exposure to the 200-ms interval, so as to promote TID learning transfer from a 100-ms interval to a 200-ms interval.

Double training improved auditory TID at a 100-ms interval by 0.27 ± 0.07 log units (t = 5.15, p < 0.001, Cohen’s d = 1.82, 95% CI [0.17, 0.38]), which was comparable to the 0.29 log unit improvement in baseline training (A100 in Fig. 1b), as well as tone frequency discrimination at a 200-ms interval by 0.18 ± 0.07 log units (t = 3.15, p = 0.004, Cohen’s d = 1.11, 95% CI [0.06,0.30]) (Fig. 2a, c). Consistent with our hypothesis, auditory TID performance was also improved at a 200-ms interval by 0.18 ± 0.03 log units (t = 3.44, p < 0.001, Cohen’s d = 1.22, 95% CI [0.08, 0.29], Fig. 2c), which was significantly higher than the corresponding 0.08 log-unit improvement in the baseline condition (A200 in Fig. 1b) (t = 2.15, p = 0.049, Cohen’s d = 1.08, 95% CI [0, 0.21]). To investigate whether the cross-interval TID learning transfer was complete, all participants continued to practice the auditory TID task at a 200-ms interval for another three daily sessions. This extra and direct training produced no further significant change of TID thresholds at a 200-ms interval (by 0.04 ± 0.03 log units; t = 0.42, p = 0.69, Cohen’d = 0.16, 95% CI [-0.09, 0.12]). Therefore, the cross-interval learning transfer had maximized auditory TID at a 200-ms interval and was complete.

**Figure 2.**
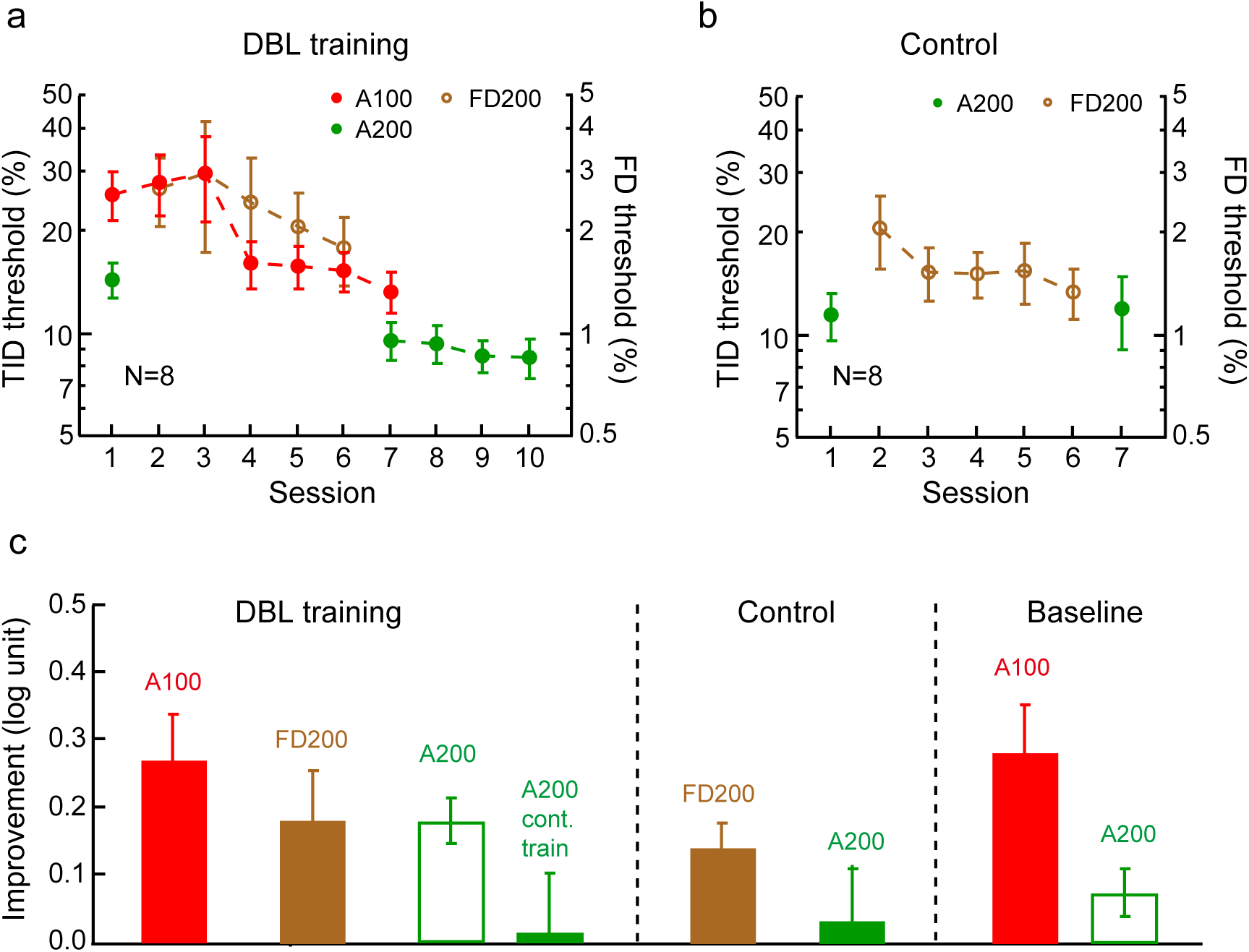
The cross-interval transfer of TID learning with double training. **a**. Double training: The mean learning curves of auditory TID at a 100-ms interval (A100, Sessions 1-7) and tone frequency discrimination task at a 200-ms interval (FD200, Sessions 2-6). Auditory TID at a 200- ms interval (A200) were tested pre- and post-double training (Sessions 1 & 7, and further practiced for 3 daily sessions (Sessions 8-10). **b**. Control: The impact of tone frequency discrimination training alone on auditory TID thresholds at the same 200-ms interval. **c**. The mean improvements of TID thresholds (A100 & A200) and tone frequency discrimination thresholds (FD200) in double training and control conditions, as well as in the earlier baseline condition (replotted from Fig. 1a).

To ensure that the learning transfer effect was not due to the tone frequency discrimination training per se, a control group only practiced tone frequency discrimination at a 200-ms interval. The training improved frequency discrimination by 0.14 ± 0.04 log units (t = 2.59, p = 0.014, Cohen’d = 0.86, 95% CI [0.03, 0.26]), but it failed to significantly change auditory TID at the same 200-ms interval (by 0.03 ± 0.08 log units; t = 0.69, p = 0.49, Cohen’s d = 0.23, 95% CI [-0.06, 0.13], Fig. 2b, c). Therefore, tone frequency discrimination was indeed orthogonal to TID, and it was the double training that enabled complete TID learning transfer across intervals.

### Complete transfer of TID learning between visual and auditory modalities with double training

Our next focus was to examine whether TID learning can transfer mutually between visual and auditory modalities with double training. Previous results only showed learning transfer from auditory to visual TID, but not in the opposite direction (Bratzke et al., 2012; McGovern et al., 2016), as we replicated earlier (Fig. 1b, d).

We first tested whether double training could enable transfer of TID learning from visual to auditory. Nine participants practiced visual TID at a 100-ms interval. They also received exposure to the auditory 100-ms interval by practicing an orthogonal tone frequency discrimination task. This double training improved visual TID at a 100 ms interval by 0.21 ± 0.03 log units (t = 4.18, p < 0.001, Cohen’s d = 1.39, 95% CI [0.11, 0.31]) and tone frequency discrimination at the same interval by 0.17 ± 0.05 log units (t = 3.13, p = 0.004, Cohen’s d = 1.04, 95% CI [0.06, 0.28]) (Fig. 3a, c). The double training also improved auditory TID at the same interval by 0.24 ± 0.04 log units (t = 4.89, p < 0.001, Cohen’s d = 1.63, 95% CI [0.14, 0.34]) (Fig. 3c). Planned comparisons suggested that this auditory TID improvement was significantly higher than that with the visual TID training baseline group (Fig. 1d) (t = 2.87, p = 0.011, Cohen’s d = 1.35, 95% CI [0.05, 0.33]). It was also not significantly different from the 0.29 log-unit improvement with direct auditory TID training with the auditory TID training baseline group (Fig. 1b) (t = 0.51, p = 0.62, Cohen’s d = 0.25, 95% CI [-0.14, 0.22]). Moreover, four participants continued to practice auditory TID for another 3 sessions, which failed to further reduce the auditory TID thresholds (by 0.02 ± 0.04 log units; t = 1.12, p = 0.34, Cohen’d = 0.56, 95% CI [-0.07, 0.15]).

**Figure 3.**
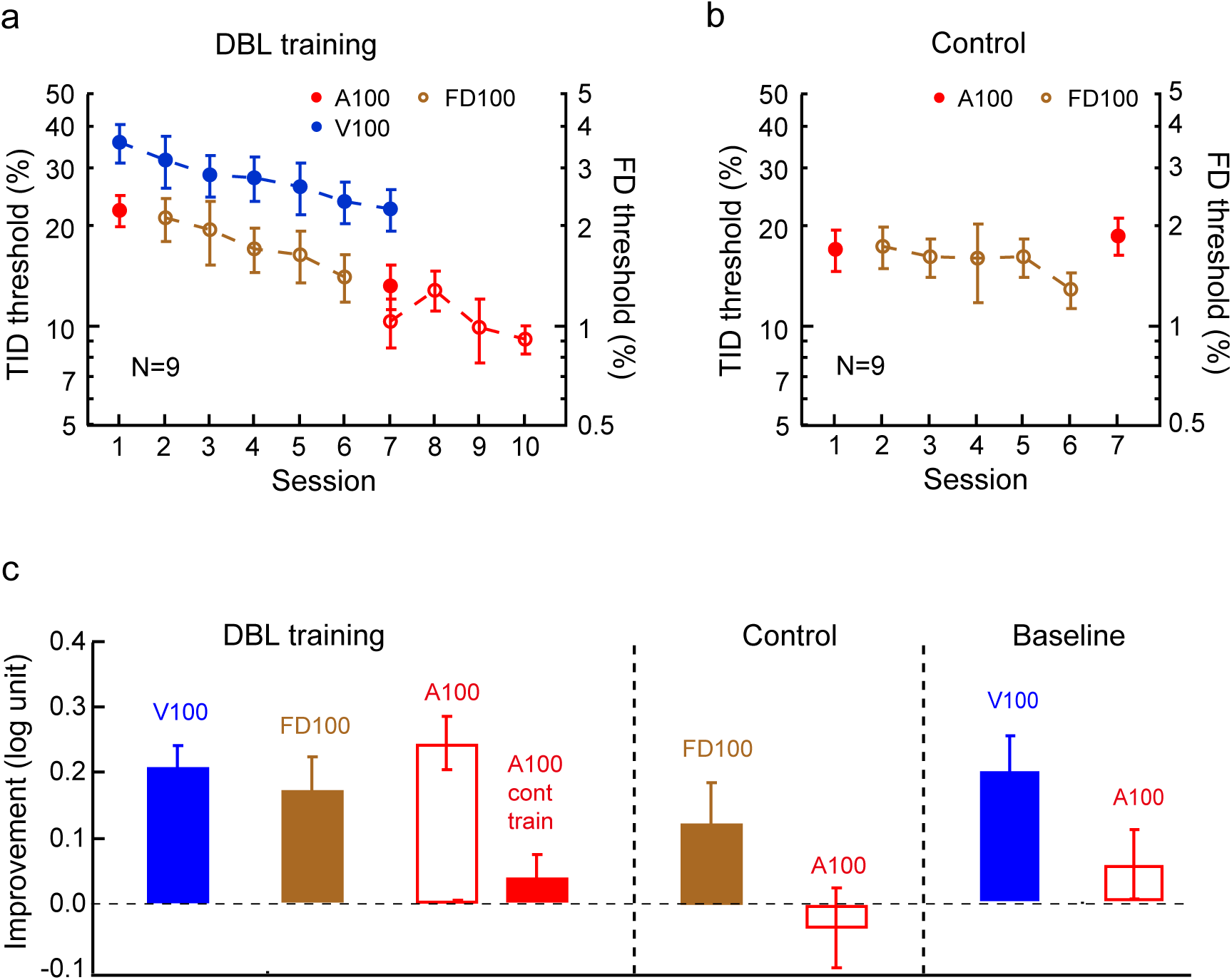
The cross-modality transfer of TID training from vision to audition with double training. **a**. Double training: The mean learning curves of visual TID at a 100-ms interval (V100, Sessions 1-7) and tone frequency discrimination task at the same interval (FD100, Sessions 2-6). Auditory TID at the same interval (A100) was tested pre- and post-double training (Sessions 1 & 7), and further practiced for 3 sessions in 4 participants (Sessions 8-10). **b**. Control: The impact of tone frequency discrimination training alone on auditory TID thresholds at the same 100-ms interval. **c**. The mean improvements of visual and auditory TID thresholds (V100 & A100) and tone frequency discrimination thresholds (FD100) in double training and control conditions, as well as in the earlier baseline condition (replotted from Fig. 1d).

To exclude the possibility that the learning transfer resulted from tone frequency discrimination training alone, we had a control group (N = 9) only practice tone frequency discrimination at a 100-ms interval. The practice improved auditory frequency discrimination by 0.12 ± 0.06 log units (t = 2.25, p = 0.031, Cohen’s d = 0.75, 95% CI [0.01, 0.24]), but it failed to make auditory TID performance at the same interval better (by -0.04 ± 0.06 log units; t = 0.74, p = 0.46, Cohen’s d = 0.25, 95% CI [-0.14, 0.06], Fig. 3b, c). Taken together, double training results and the control data suggested that double training enabled full learning transfer from visual TID to auditory TID, in contrast to insignificant transfer in the baseline condition (Fig. 1c, d).

We were also curious whether double training could lead to better auditory-to-visual TID learning transfer than the baseline auditory TID training only condition, in which the visual TID improvement from auditory TID training was about 70% of that from direct visual TID training (Fig. 1b, d). Eight new participants practiced auditory TID and visual contrast discrimination, both at a 100-ms interval, in alternating blocks of trials in the same training sessions. Training improved auditory TID threshold by 0.29 ± 0.04 log units (t = 5.41, p < 0.001, Cohen’s d = 1.91, 95% CI [0.18, 0.39]) and visual contrast discrimination by 0.37 ± 0.20 in d’ (t = 2.06, p = 0.058, Cohen’s d = 0.73, 95% CI [-0.01, 0.74]), as well as visual TID at the same 100-ms interval by 0.26 ± 0.03 log units (t = 4.95, p < 0.001, Cohen’s d = 1.75, 95% CI [0.16, 0.37]) (Fig. 4a, c). Planned comparisons showed that the visual TID improvement was significantly higher than the 0.14 log-unit improvement in the auditory TID training baseline group (Fig. 1b) (t = 2.51, p = 0.026, Cohen’s d = 1.25, 95% CI [0.02, 0.24]), but it did not differ significantly from the 0.20 log-unit improvement through direct visual TID training in the visual TID training baseline group (Fig. 1c) (t = 1.06, p = 0.31, Cohen’s d = 0.52, 95% CI [-0.07, 0.20]). Four participants continued to practice visual TID for another 3 sessions, which failed to significantly change the visual TID thresholds (by -0.07 ± 0.10 log units; t = 0.72, p = 0.53, Cohen’d = 0.36, 95% CI [-0.25, 0.40]), suggesting that the visual TID performance had maximized after double training.

**Figure 4.**
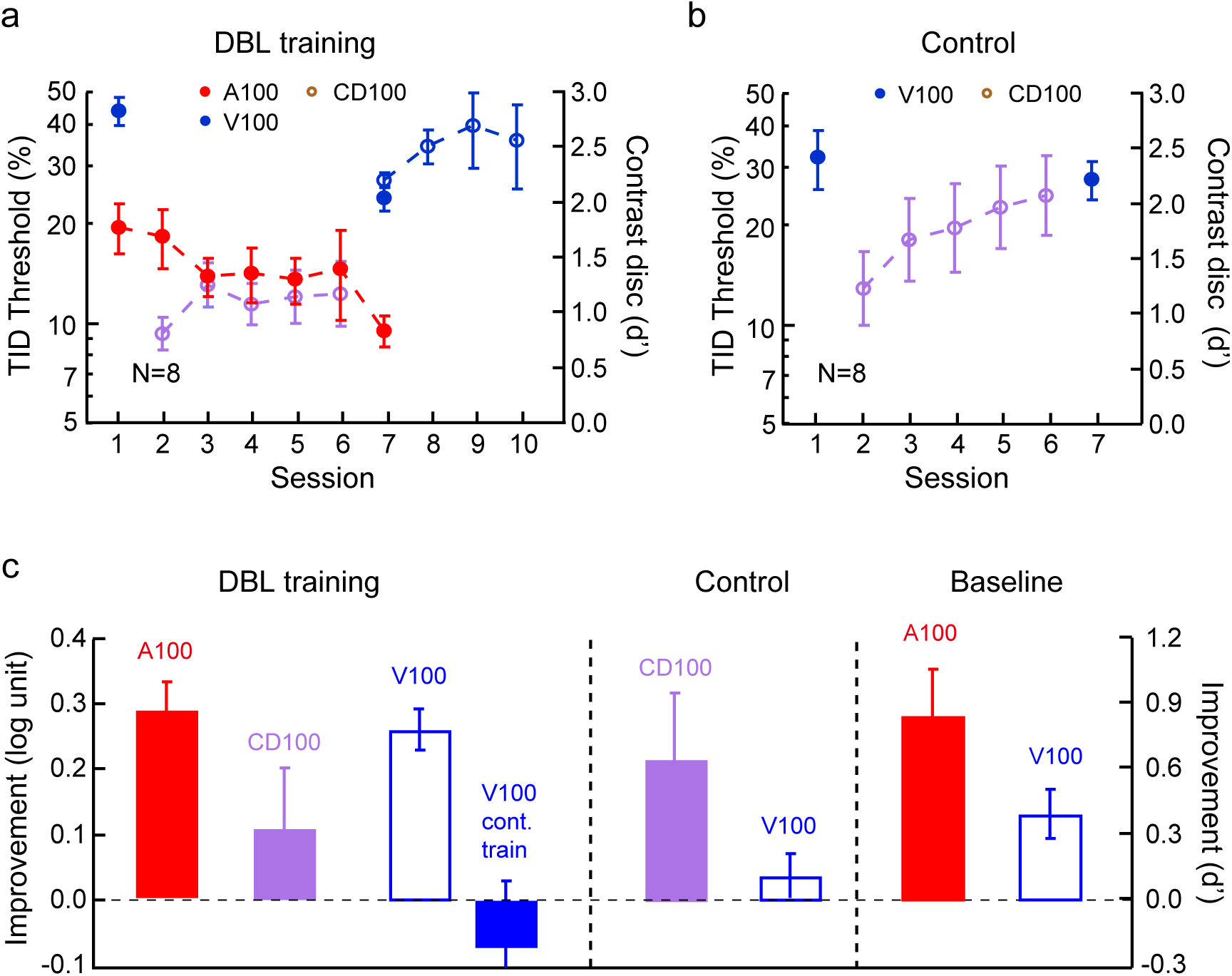
The cross-modality transfer of TID training from auditory to visual with double training. **a**. Double training: The mean learning curves of auditory TID at a 100-ms interval (A100, Sessions 1-7) and visual contrast discrimination at the same interval (CD100, Sessions 2- 6). Visual TID at the same interval (A100) was tested pre- and post-double training (Sessions 1 & 7), and further practiced for 3 sessions in 4 participants (Sessions 8-10). **b**. Control: The impact of visual contrast discrimination training alone on visual TID thresholds at the same 100-ms interval. **c**. The mean improvements of auditory and visual TID thresholds (A100 & V100) and visual contrast discrimination thresholds (CD100) in double training and control conditions, as well as in the earlier baseline condition (replotted from Fig. 1b).

Again a control group of participants (N = 8) practiced visual contrast discrimination only, which improved contrast thresholds by 0.84 ± 0.15 in d’ (t = 4.74, p < 0.001, Cohen’s d = 1.68, 95% CI [0.46, 1.22]). But this practice had no significant impact on visual TID at the same 100-ms interval (by 0.04 ± 0.03 log units; t = 0.73, p = 0.47, Cohen’s d = 0.26, 95% CI [-0.07, 0.14], Fig. 4b & c). These and above double training results thus demonstrated complete auditory-to-visual TID learning transfer with double training.

## Discussion

In this study we demonstrated with a double training technique that TID learning can transfer completely to a new interval, and mutually completely between visual and auditory modalities. These new findings contradict previous observations of interval specificity and asymmetric auditory-to-visual transfer of TID learning through conventional training. When the major constraints on interpreting subsecond timing by these observations are no longer in place, a new account that unifies the long-debated models of dedicated and distributed mechanisms of subsecond timing becomes possible.

The full transfer of TID learning from a 100-ms to a 200-ms interval suggests an interval-invariant abstract representation of subsecond time in the brain. This high-order time representation is reminiscent of the idea of a dedicated centralized clock. Models of a dedicated centralized clock have been challenged because they have difficulty to account for the interval specificity in TID learning (Bueti & Buonomano, 2014). In addition, although spatial transfers of TID learning at the same interval, like from a trained tone frequency to a new frequency (Wright et al., 1997), may seemingly support a common dedicated timing mechanism, it could alternatively result from a more precise representation of a specific interval in the working memory (Bueti & Buonomano, 2014). However, an interval invariant representation of subsecond time is consistent with the idea of a centralized clock (Creelman, 1962; Treisman, 1963), especially if we take the original centralized clock models as assumptions for some centralized timing mechanisms that can process multiple durations/intervals, without caring too much about these early models’ specifics. These centralized mechanisms or the clock may represent different durations/intervals in an abstract manner (e.g., through standardization of incoming neural inputs representing different durations/intervals). Moreover, perceptual learning may improve the precision of this abstract time representation, so that TID learning can transfer from a trained interval to a new interval.

The asymmetric transfer of TID learning from audition to vision has been interpreted as auditory timing mechanisms, which may have less internal noise and thus be more precise, dominating the TID process (Bratzke et al., 2012; McGovern et al., 2016). Parallel to this asymmetric transfer is the observations that visual perceptual learning at zero or low external noise can transfer to high external noise, but not vice versa (Dosher & Lu, 2005). Also parallel to the current double training results is our recent report of mutual complete learning transfer of visual learning between zero and high external noise levels with double training (Xie & Yu, 2019). Since the initial asymmetric transfer with conventional training is inconsistent with an improved working memory account (Bueti & Buonomano, 2014), the mutual transfers of TID learning between visual and auditory modalities would add evidence for an abstract representation of subsecond time, which is independent of not only specific intervals, but also sensory modalities and timing precisions.

It is at least conceptually feasible to come up with a unified model of subsecond timing. A subsecond duration/interval is first responded by various distributed neural mechanisms, such as temporal ramps in neuronal firing (Durstewitz, 2003) or peaks in sequential activations of neuron pools (Hass et al., 2008). The exact duration/interval is then readout from responding neurons or circuities, following certain decision rules. In the case the same time information is picked up by multiple mechanisms or sensory modalities, the information from multiple sources may be integrated on the basis of Bayesian priors (e.g., more precise the mechanisms or modalities are, more weight is assigned to the time information they carry in the integration equation) (Cicchini et al., 2012; Shi et al., 2013; Hass & Durstewitz, 2016). On top of these duration/interval-specific processes, there is also an abstract duration/interval-invariant representation of subsecond time, as our double training results suggest, which may serve as a master clock. Merchant et al. (2013) once reported modality invariant (visual & auditory) but interval specific time representation in macaque medial premotor cortex. We thus suspect that a modality and interval invariant time representation may occur in brain areas further downstream. In the context of such as unified framework, training may improve the precision of not only duration/interval-specific mechanisms, but also the duration/interval-invariant master clock, so that learning can transfer from a trained interval to a new interval.

It remains a mystery why TID learning is interval specific in the first place with conventional training. Our general understanding on this matter mainly comes from visual perceptual learning studies of ours. After demonstrating learning transfer to new retinal locations and orientations (Xiao et al., 2008; Zhang et al., 2010), we further discovered that stimulation of the transfer location or orientation through either bottom-up exposure or top-down attention is sufficient to induce learning transfer (Xiong et al., 2016). We also found that significant occipital P1-N1 changes are associated with learning transfer to a new location, but are absent when learning is location specific (Zhang et al., 2013). Therefore, we propose that sensory features such as stimulus orientation (or time interval in the current case) have abstract or conceptual representations in the brain. Moreover, perceptual learning operates at a conceptual level (i.e., reweighting standardized sensory inputs) (Wang et al., 2016), so that learning can in principle be applied to new sensory inputs to enable transfer. However, whether learning can actually transfer depends on the functional connections between high-level learning and new sensory puts that are unattended and even possibly suppressed during training. Such functional connections can be established through exposure to new sensory inputs in double training, so that learning becomes transferrable. As for the non-transfer of learning from visual to auditory TID, we assume that while participants have learned an abstract subsecond time representation from noisier visual interval signals, they may also need to learn the less noisy auditory interval signals through tone frequency discrimination at the same physical interval. After that TID learning transfers from vision to audition.

## Methods

### Participants and Apparatus

Data were collected from 68 college students (46 female, 20.9±2.2 years old) randomly allocated to each experiment. All of them had normal or corrected-to-normal vision and normal hearing (pure-tone thresholds ≤ 20 dB hearing level across 0.5–6 kHz). They had no previous experience with visual psychophysical or psychoacoustic experiments, and no knowledge of the purpose of the study. Informed consent was obtained from each participant prior to data collection. The study was approved by the Peking University IRB.

Experiments were run in an anechoic booth. The stimuli were generated with a Matlab-based Psychtoolbox software (Pelli, 1997). Auditory stimuli were diotic, presented by a pair of Sennheiser HD-499 headphones. Visual stimuli were presented on a 19-inch Sony G420 CRT monitor with a resolution of 800 pixel × 600 pixel and a refresh rate of 160 Hz. The luminance of the monitor was linearized by an 8-bit look-up table, with a mean luminance of 43.5 cd/m^2^. A chin-and-head rest stabilized the head of the observer.

### Stimuli and Procedures

The auditory stimuli were two 15-ms tone pips separated by a 100 ms or 200 ms interval. Each tone contained a 5-ms cosine ramp at each end, and was fixed at 1 kHz and 86 dB SPL. The visual stimuli were two 15-ms Gabor gratings, separated by a 100 ms interval. Each Gabor had a fixed orientation (0°), spatial frequency (1 cycle/deg), and contrast (100%). The length of the interval was the difference between the offset of the first stimulus and the onset of the second stimulus.

The temporal interval discrimination thresholds were measured with a method of constant stimuli. In each forced-choice trial, a visual fixation was first centered on the computer screen for 300 ms, then two pairs of stimuli, one with a standard interval (t) and the other with a comparison interval (t + Δt), were subsequently presented in a random order with a 900-ms gap in between. The standard interval (t) were 100 or 200 ms. The participants pressed the left or right arrow to indicate whether the first or the second pair of stimuli had a longer interval. A happy or sad cartoon face was shown after each response to indicate a correct or wrong response. A blank screen was presented before the next trial for a random duration (500-1000ms). The Δt was set at 6 levels for each condition (A100: ±20.1, ±13.4, ±6.7 ms; V100: ±33.5, ±20.1, ±6.7 ms; A200: ±40.1, ±26.8, ±13.4 ms; V200: ±67.0, ±40.1, ±13.4 ms), and was adjusted individually if necessary to ensure a sufficient range of correct rates. Each level was repeated 10 times in a block of 60 trials, for a total of 5 blocks.

The psychometric function was fitted with 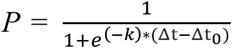, where *P* was the rate of reporting the comparison interval being longer at each Δt, k was the slope, and Δt_0_ was the point of subjective equivalence. The temporal interval discrimination threshold was estimated as being half the interquartile range of the function: Threshold = (Δt_.75_- Δt_.25_)/2.

The stimuli for tone frequency discrimination were the same as those for auditory temporal interval discrimination, excepted that the frequencies of two pairs of pips were changed while the temporal intervals were fixed. The two pairs of pips, one pair at a standard frequency of 1 kHz and the other at a higher comparison frequency (1 kHz + Δf), were presented subsequently in a random order in each trial. The participants pressed the left or right arrows to indicate whether the first or second pair of tone pips had a higher frequency. Happy or sad cartoon face was provided as feedback.

The frequency discrimination thresholds were measured with a 2IFC staircase procedure. The starting frequency difference (Δf) between the standard and comparison stimuli was 50%, which decreased by a factor of 2 after every correct response until the first incorrect response. After that the Δf was varied by a factor of 1.414 following a 3-down-1-up rule for a 79% correct rate. Each staircase ended after 60 trials. The threshold was calculated as the mean of the last 40 trials.

The stimuli used for visual contrast discrimination was the same as those for visual temporal interval discrimination, except that the Gabor contrast was varied while the interval was fixed (100 ms). Only one pair of Gabor was presented in each trial. In 80% of the trials, the two Gabors had identical contrast, which randomized from 0.15 to 1. In the remaining 20% trials, the contrasts of the two Gabors differed by 50%. The participants judged whether the two Gabors had identical contrast. Happy or sad cartoon face was provided as feedback. The d’ values were calculated to measure the contrast discrimination performance.

Each experiment consisted of a pre-training session, five training sessions, and a post-training session on separate days. The experiment was completed within 7-13 days, with inter-session gaps of no more than 2 days. Each single-training session consisted of 16 blocks of trials and lasted for approximately 1.5 hours. Each double-training session consisted of 10 blocks of trials for the primary task and 10 blocks of trials for the secondary task in an alternating order, and lasted for approximately 2 hours.

### Sample Size

The sample size was decided on the basis of a previous temporal interval discrimination learning study that used similar stimuli (Fig. 4, 100 ms – 1 kHz condition in Wright et al. (1997)). In our study, learning and transfer involved comparisons between pre- to post-training thresholds in all experiments. To achieve 80% power at p = 0.05, for a similar effect size of Cohen’s d = 1.34 obtained from Wright et al. when comparing pre- and post-training thresholds, a sample size of 7 would be required. We used a sample size of 9 for each experiment, including dropouts of participants.

### Data analysis

Data were analyzed using the R software (R_Core_Team, 2015). The primary analysis was performed by Linear Mixed Effects (LME) modeling to examine the training and transfer effects of the TID task, using the “lmer” function from the “lme4” package (Pinheiro & Bates, 2000). All groups’ data were included in a single LME model to reduce Type-I error. The TID thresholds were first log-transformed to achieve normal distributions (Shapiro-Wilk test before log-transformation: p < 0.001 for auditory TID thresholds at 100 ms and 200 ms intervals, and visual TID thresholds at 100 ms intervals; Shapiro-Wilk test after log-transformation: p = 0.28, 0.76, and 0.60 for corresponding TID thresholds). Specifically, the models treated Threshold as the dependent variable, Condition (training and transfer conditions), Test (pre- & post-training tests), and Group as the fixed effects. For each participant, we included random slopes for Test and Condition and conducted model selection based on their significance. The significance of the fixed effects was evaluated by the anova function in the “lmerTest”. The significance of the random-effect components was evaluated by the Likelihood-ratio test. Post-hoc analyses were then conducted on the best fitting model to test the learning and transfer effects under each condition in each group. The Post-hoc analysis was conducted by the “emmeans” package (Piepho, 2004).

The LME analysis revealed significant main effect of Test (F(1, 55) = 68.71, p < 0.001) and Condition (F(2,28) = 86.72, p < 0.001), but not Group (F(7,44) = 0.18, p = 0.99). There were significant interactions between Group and Test (F(7,57) = 5.43, p < 0.001), between Test and Condition (F(2,41) = 5.41, p = 0.008) and a three-way interaction between Group, Test and Condition (F(4,41) = 4.16, p = 0.006). The post-hoc analysis of the learning and transfer effects were reported with relevant results.

## Acknowledgements

This research was supported by a Natural Science Foundation of China grant 31230030 and funds from Center for Life Sciences, Peking University. We thank the comments from Dean Buonomano.

